# Beyond the Hypercube: Evolutionary Accessibility of Fitness Landscapes with Realistic Mutational Networks

**DOI:** 10.1101/067819

**Authors:** Marcin Zagorski, Zdzislaw Burda, Bartlomiej Waclaw

## Abstract

Evolutionary pathways describe trajectories of biological evolution in the space of different variants of organisms (genotypes). The probability of existence and the number of evolutionary pathways that lead from a given genotype to a better-adapted genotype are important measures of accessibility of local fitness optima and the reproducibility of evolution. Both quantities have been studied in simple mathematical models where genotypes are represented as binary sequences of two types of basic units, and the network of permitted mutations between the genotypes is a hypercube graph. However, it is unclear how these results translate to the biologically relevant case in which genotypes are represented by sequences of more than two units, for example four nucleotides (DNA) or 20 aminoacids (proteins), and the mutational graph is not the hypercube. Here we investigate accessibility of the best-adapted genotype in the general case of *K* > 2 units. Using computer generated and experimental fitness landscapes we show that accessibility of the global fitness maximum increases with *K* and can be much higher than for binary sequences. The increase in accessibility comes from the increase in the number of indirect trajectories exploited by evolution for higher *K*. As one of the consequences, the fraction of genotypes that are accessible increases by three orders of magnitude when the number of units K increases from 2 to 16 for landscapes of size *N* ~ 10^6^ genotypes. This suggests that evolution can follow many different trajectories on such landscapes and the reconstruction of evolutionary pathways from experimental data might be an extremely difficult task.

## I. INTRODUCTION

Biological evolution can be visualised as a dynamical process in which reproduction and mutations cause organisms to move in the “fitness landscape” - a complex, multidimensional landscape full of ridges and local peaks in which space represents genetic variants (genotypes) of organisms, and “height” corresponds to organism’s fitness - a measure of the reproductive success.

The structure of the fitness landscape (FL) affects the speed and predictability of biological evolution and is thus very important for practical reasons such as forecasting the evolution of resistance to antibiotics [1–6]. However, fitness landscapes are astronomically large even for the simplest organisms: the number of possible genotypes with the same genome size as the smallest known genome to-date *(Carsonella ruddii* [7], *L* = 160kbp) is *N* = 4^160000^ ≈ 10^96329^ genotypes. For this reason only small fragments of fitness landscapes have been determined experimentally [2, 8–12] and progress has been made mostly by studying adaptation in computer-generated landscapes (see e.g. [13–15]) or in simple theoretical models (see e.g. House-of-Cards [16], NK [17], Mount Fuji [18], holey landscapes [19]). In these models genotypes are typically represented as binary sequences, where 0 and 1 correspond to two different alleles of a gene or a particular point mutation being absent/present. An additional assumption that offspring differ from their parent by only one character in the sequence causes the graph of permitted mutations to be a hypercube graph, and this greatly facilitates mathematical analysis [20, 21].

However, the space of real genotypes and their mutational network differ significantly from the space of binary sequences and the hypercube graph. A DNA sequence is made of four different nucleotides (A,T,G,C), hence at the most fundamental level genotypes must be represented as sequences of *K* = 4 different characters. On a more coarse-grained level, assuming that fitness is determined primarily by protein structure, a genotype can be represented as a sequence of characters from a set of *K* > 20 symbols which correspond to 20 standard amino acids [22, 23] and their post-translational modifications. Finally, if we consider genotypes as sequences of genes, *K* is equal to the number of different variants (alleles) of genes and is practically infinite. All these examples are valid descriptions of the genotype space but they lead to different structures of the corresponding fitness landscape: different genotype-to-fitness mappings and different structures of the graphs of possible transitions (mutations) between the genotypes.

Here we investigate how the global structure of the fitness landscape is affected by the number *K* of different “units”, or characters, of which sequences representing genotypes are made, and the structure of the corresponding mutational graph (Fig. 1). We focus on evolutionary accessibility of the global fitness maximum. An accessible pathway is a trajectory in the genotype space along which fitness increases monotonically. Accessible pathways are important because in the low mutation-strong selection limit [24–26] such pathways are good approximations to evolutionary trajectories obtained in computer models of biological evolution, regardless of the details of the model. Accessible pathways are thus expected to represent real evolutionary pathways, at least in some experiments [2].

**Fig 1.**
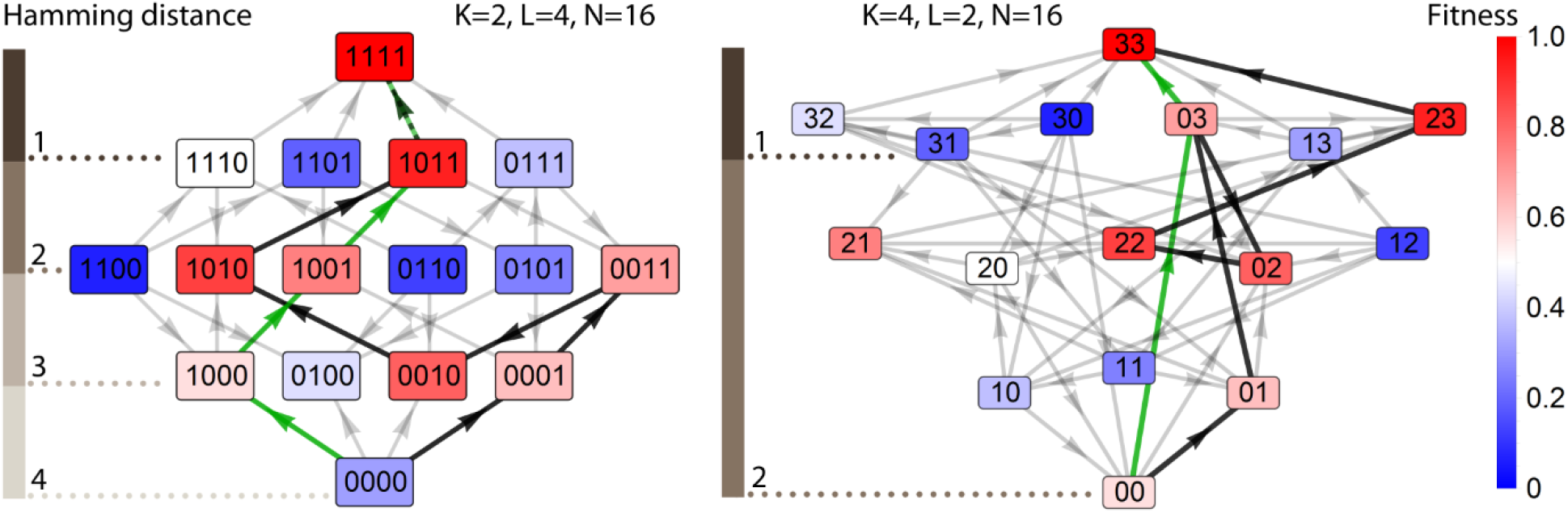
Different numbers of sequence-coding units *K* result in different mutational structures of the fitness landscape. Examples of two fitness landscapes with *K* = 2 and *K* = 4 and the same number of genotypes *N* = 16. Grey arrows show the direction of increasing fitness. Two accessible pathways (black = with indirect mutations, green = without indirect mutations) have been highlighted. For *K* = 4, indirect mutations can be either backward (increasing the Hamming distance from the target genotype) or distance-neutral (the Hamming distance remains unchanged).

While we intuitively expect that increasing the number of coding units and hence the connectivity of the mutational graph should increase accessibility if the total number of genotypes *N* = *K^L^* is kept fixed, it is not obvious how increasing *L* affects accessibility for different *K*. In the case of well-studied binary landscapes (*K* = 2) the probability that a maximally random (House-of-Cards) fitness landscape has at least one accessible pathway approaches zero as *L* → ∞ [26–28] but, to our knowledge, the case *K* > 2 has not been studied. It is also unclear what happens for *K* > 2 in the presence of correlations between the fitnesses that exist in many real fitness landscapes [9].

To address these questions, we analyse computer-generated and experimental fitness landscapes with various *K* and *L*. First, we show that if maximally random landscapes with the same number of genotypes *N* = *K^L^* but different *K* are compared, larger *K* leads to higher accessibility. In contrast to the binary case *K* = 2, in the large-*L* limit the average landscape with *K* > 2 contains at least one accessible pathway. We demonstrate that as *K* increases, accessible pathways become longer and contain an increasing number of indirect mutations which do not decrease the distance to the fittest genotype. We also show that accessibility depends strongly on the fitness of the initial genotype, *f*_0_, and that a transition occurs at some critical *f*_0_ above which there are no accessible pathways. Finally, we confirm our predictions for two experimentally determined fitness landscapes.

### Modelling framework

We consider the space of all possible sequences of length *L* made of symbols taken from a *K*-letter alphabet. Each sequence *i* = 0, …, *K^L^* – 1 represents a genotype in the discrete genotype space, with *i* = 0 corresponding to sequence {0, 0, 0, …, 0} and *i* = *K^L^* – 1 to {*K* – 1, *K* – 1, *K* – 1, …, *K*–1}. The genotypes are connected by a network of mutations that change one genotype into another one. We assume that any position in the sequence can mutate with equal probability and that a mutation at a given position changes the symbol at this position to another randomly selected symbol. Therefore, two genotypes are connected if they differ at exactly one position.

To create a fitness landscape, we assign random fitnesses {*f_i_*} to all genotypes. Each fitness *f_i_* is a random number drawn independently from the probability distribution uniform on [0…1). Next, we re-index all genotypes such that genotype {*K* – 1, *K* – 1, *K* – 1, …, *K* – 1} has the maximum fitness (Methods). This procedure creates an ensemble of maximally-random or “rugged” fitness landscapes with no correlations between the fitnesses of adjacent genotypes. This is arguably the simplest non-trivial model of the fitness landscape with many local maxima and minima and thus it presents an ideal test ground to investigate the role of connectivity on the accessibility of FLs. Although real landscapes are known to be moderately correlated [9, 29–31], we shall show later that our conclusions are robust in the presence of such correlations.

We define a pathway in the genotype space as a trajectory that connects two given genotypes following the links of the mutational graph. We call a pathway accessible if fitness increases monotonically from the start to the end point along the pathway. We consider only such evolutionary trajectories that begin at genotype {0, 0,0, …, 0} and end at the best-fit, antipodal genotype {*K* – 1, *K* – 1, *K* – 1, …,*K* – 1}. Following Ref. [9], we call a FL accessible if there is at least one accessible path between the target and the initial genotype (Fig. 1). We also define accessibility A as the probability that a FL randomly selected from our statistical ensemble is accessible.

## II. RESULTS

### Computational model

#### Accessibility increases with K

We investigated how accessibility varied with the number of genotypes *N* = *K^L^* for different *K*. We first considered only pathways without “backward mutations”, that is pathways along which the Hamming distance from the initial genotype increased (or remained the same for *K* > 2) with each step. Figure 2a shows that accessibility A slowly decreases with increasing *N*, for all *K*. When *K* = 2, we recover the mathematical result of Hegarty et al. [27] that A tends to zero for *N* → ∞. However, when *K* > 2, accessibility approaches a finite, non-zero value in the large-*N* limit. The estimated asymptotic accessibility *A*_∞_ = *A*(*N* → ∞) increases with *K* and equals 47%, 69% and 81% for *K* = 4, 8, and 16 (Fig. S1) (standard errors < 1%). Therefore, for sufficiently long sequences accessibility is larger for a larger number of basic unit types.

**Fig 2.**
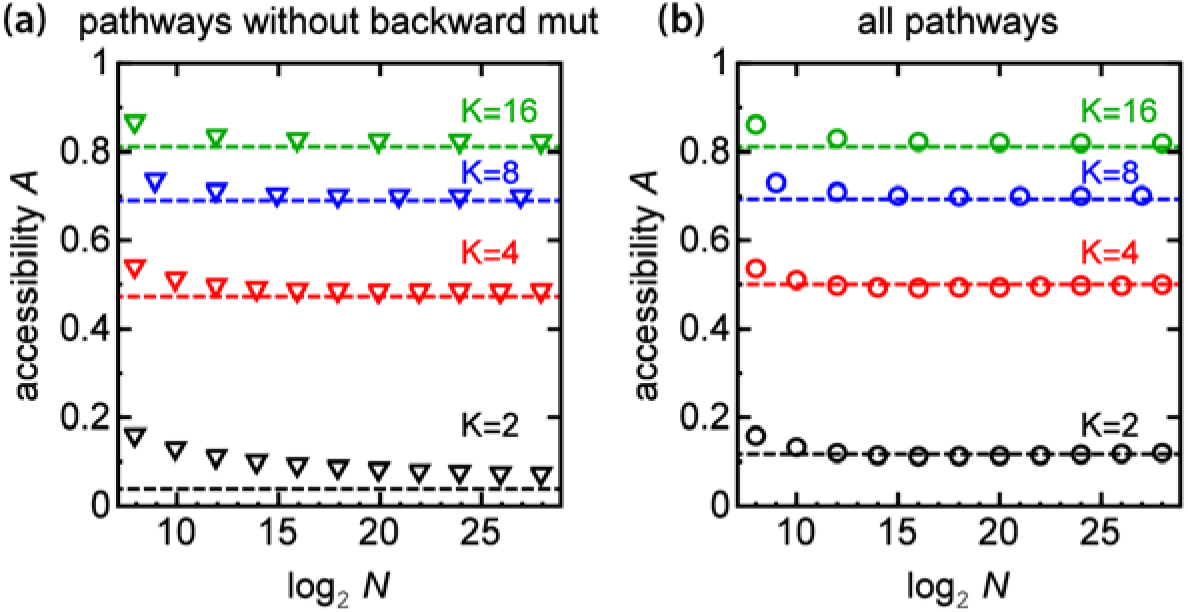
Accessibility increases with increasing number of unit types *K*. **(a)** Plots of accessibility *A* versus the number of genotypes *N* for different *K* (black, red, blue, and green triangles) and for pathways without backward mutations. *A* tends to zero as *N* → ∞ for *K*=2. **(b)** Accessibility versus *N* for all pathways, including indirect ones (circles). In both panels the dashed lines correspond to asymptotic estimates of accessibility *A*_∞_ (cf. Fig. S1).

Figure 2b shows that the same trend holds when all pathways and not just the ones without backward mutations are considered. However, in the latter case *A_∞_* remains finite also for *K* = 2. A comparison of panels (a),(b) in Fig. 2 shows that, except for the case *K* = 2, accessibility is almost identical regardless of whether pathways with backward mutations are included or not. When *K* = 2 we recover the analytical prediction *A* = 0.1186 … from Berestycki et al. [28]. We also note that if only shortest pathways are included for *K* > 2, accessibility tends to zero as *N* → ∞ because in this case only mutations that change the symbol “0” to *K* – 1 are permitted and thus the space of permitted genotypes is restricted to that of binary sequences (*K* = 2). Since pathways without backward mutations form only a small subset of all pathways for *K* > 2 (Fig. S2) and hence they are less likely to be selected by evolution in the presence of a large number of other pathways with backward mutations, in the rest of the paper we shall consider all pathways when discussing accessibility.

#### Larger K increases the length and the fraction of indirect mutations in accessible pathways

We next investigated the attributes of individual accessible pathways: their length and the fraction of backward mutations. Increasing *K* while keeping *L* fixed makes accessible pathways longer (Fig. 3a). Interestingly, the average length scales as ~ 0.5*LK* and only weakly depends on *N* (Fig. S3). Notice that by fixing *L* we keep the same mutational (Hamming) distance between the initial and target genotype, thus any increase in pathway length for *K* > 2 must be caused by mutations that either do not change the Hamming distance from the initial genotype (distance-neutral mutations) or decrease it (backward mutations), which we jointly refer to as indirect mutations. This is indeed what happens; Fig. S2 shows that the probability that an accessible path has backward or distance-neutral mutations increases with *K* if we compare fitness landscapes with the same *L*. Moreover, the higher *K* the more dominating are distance-neutral mutations (Fig. 3a). For example, for *K* = 16 accessible pathways have on average 5% of backward, 15% of forward and 80% of distance-neutral mutations. This is in contrast to the binary case *K* = 2 in which forward mutations account for 98% of all mutations, and the remaining 2% are backward mutations. Keeping *N* or *K* fixed while changing *K* or *L* (Fig. 3b,c) does not qualitatively change this picture. Hence, higher accessibility of FLs with multiple coding units comes with a price: accessible pathways become longer, and more mutational steps are required to reach the best-fit genotype than in the binary case (*K* = 2).

**Fig 3.**
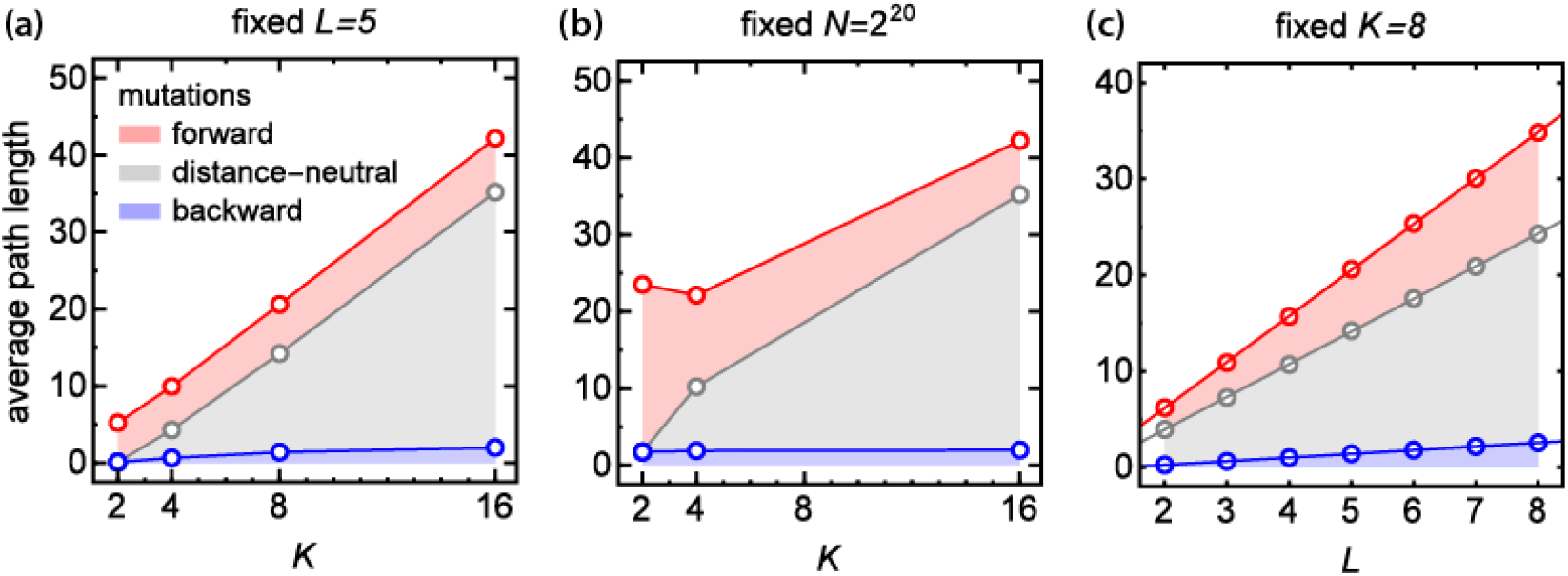
Indirect mutations dominate evolutionary trajectories for fitness landscapes with *K >* 2. **(a)** Average length of accessible pathway for a fixed length of genotype *L*. Fractions of forward, distance-neutral, and backward mutations are indicated by coloured areas between the curves, for example the fraction of forward mutations corresponds to the distance between the red and the gray curve. **(b)** Average length of accessible pathway for a fixed number of genotypes *N*. **(c)** Average length of accessible pathway for a fixed number of coding units *K* (circles) with straight lines fitted to the data (lines).

#### Initial fitness affects the number of pathways and accessibility

So far we have been assigning a random fitness to the initial genotype. We may however expect that the fitness *f*_0_ of the initial genotype strongly affects the number of accessible pathways from that genotype to the antipodal, best-fit genotype. If *f*_0_ was close to the maximal fitness we would intuitively expect few (or none) pathways, whereas for small *f*_0_ such pathways would be more likely. This has been indeed observed for binary sequences (*K* = 2) [27, 28, 32]. Figure 4a shows that this is also the case for *K* > 2: the number of accessible pathways decreases with increasing *f*_0_. Moreover, accessibility sharply decreases from ≈ 1 to zero as *f*_0_ crosses a critical *f*_crit_, different for each *K* (Fig. 4b). This transition between accessible and inaccessible FLs becomes sharper with increasing *L*, and for *L* → ∞ the numerical value of *f*_crit_ approaches *A*_∞_. Thus, for *L* large enough, while accessible pathways exist for *K* = 2 only if the initial fitness is very low (*f*_0_ < 0.1), the number of such pathways is non-zero even for relatively large *f*_0_ when *K* > 2.

**Fig 4.**
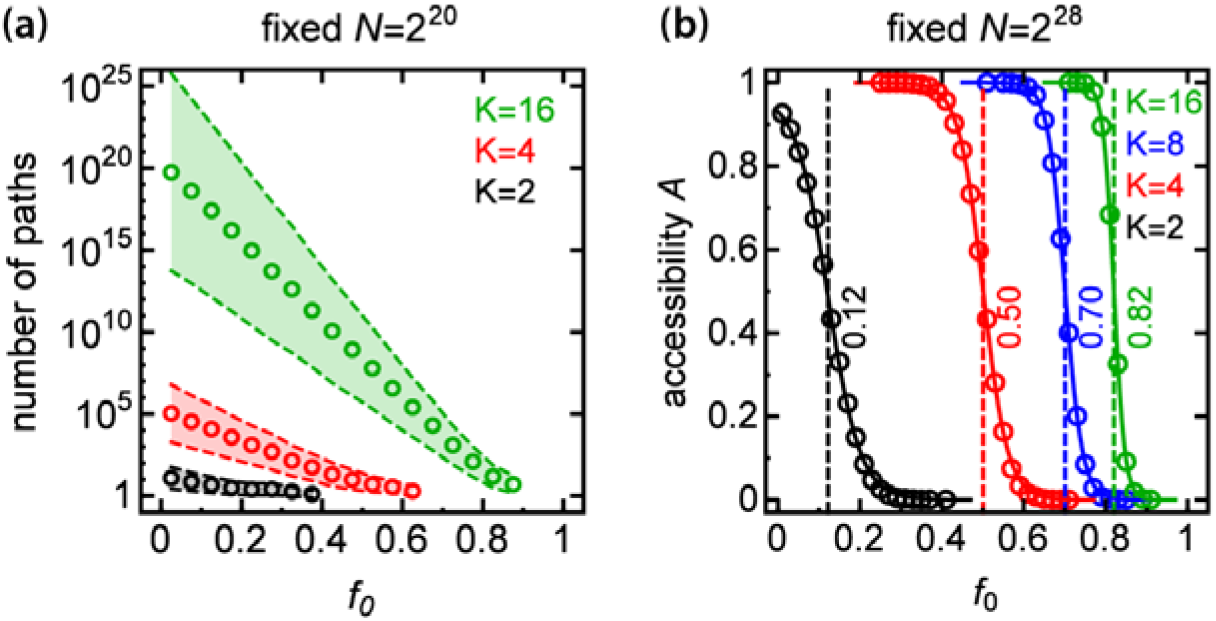
Accessibility strongly depends on the initial fitness. **(a)** Average number of accessible pathways as a function of initial fitness *f*_0_ for different *K* and for *N* = 2^20^. Shaded areas represent standard errors. **(b)** Accessibility *A* as a function of initial fitness *f*_0_ for different *K* and for *N* = 2^28^. A steep decrease in accessibility at some critical *f*_0_ (dashed line) indicates that the probability of reaching the best-fit genotype can be very sensitive to the initial fitness.

#### Accessible pathways can cover most of the FL

Figure 4a shows that the number of accessible pathways increases by several orders of magnitude when *K* increases from 2 to 16. This is caused by a rapid increase in the number of genotypes belonging to accessible pathways, i.e., the coverage of the FL (Fig. 5). In particular, for *K* = 2 the average coverage of the FL equals 0.02%, whereas for *K* = 16 around 25% of genotypes belong to accessible pathways. Coverage can exceed 50% when the initial fitness *f*_0_ is small (Fig. S4). The increase in the number of accessible pathways and accessible genotypes has important consequences for the *a posteriori* predictability of biological evolution: given the initial and final genotypes, we cannot reliably predict the pathway biological evolution might have followed if *K* > 2.

**Fig 5.**
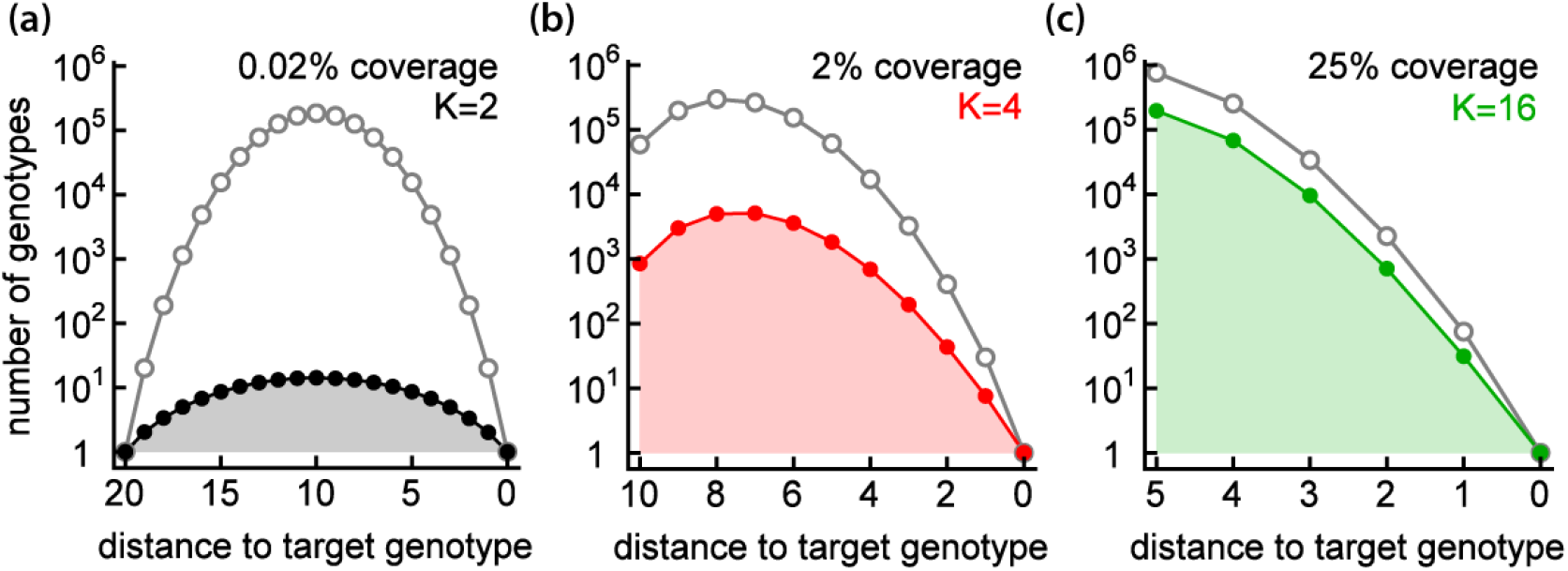
Accessible pathways cover a large part of FL for *K*>2. The total number of genotypes (grey) and the genotypes belonging to accessible pathways (colours black, red, and green) as a function of Hamming distance from the target genotype, for fitness landscapes with different number of coding units *K* = 2,4,16 (panels a,b,c) and the same number of genotypes *N* = 2^20^. The shaded area under the curve corresponds to the total number of genotypes (for any distance) that belong to accessible pathways. Coverage (the fraction of genotypes in accessible pathways) is also shown.

### Experimental fitness landscapes

We next verified the predictions of our computer model using two experimentally determined fitness landscapes. We first analysed the landscape of DNA-protein affinities [31], with binding affinity to the fluorescent protein allophycocyanin as a proxy for fitness. By selecting subsets of genotypes (Methods) we generated two ensembles of fitness landscapes with *L* = 5, *K* = 4 and *L* = 10, *K* = 2 (*N* = 1024 genotypes in each case).

We obtained accessibility *A* = 0.669(5) for *K* = 2 and *A* = 0.864(4) for *K* = 4, thus confirming our prediction that *A* increases with *K* (Fig. 6a-c). The values of *A* are higher than for the maximally random landscape (Fig. 2) because the fitness values are negatively correlated with the distance to the target genotype, i.e., the closer to the target genotype, the higher is the fitness (Fig. 6b). If we randomize the fitness landscapes by randomly swapping the fitnesses, the expected A decreases and it matches the value obtained for the maximally random landscape. Correlations between the fitness and the distance of a genotype to the target genotype can therefore significantly increase accessibility. We note that this is not caused by the difference between the distributions of *f*_0_ for randomized and original FLs because these distributions strongly overlap (Fig. 6c). This observation indicates that accessibility can be enhanced by fitness-to-distance correlations in FLs, in accordance with [26].

**Fig 6.**
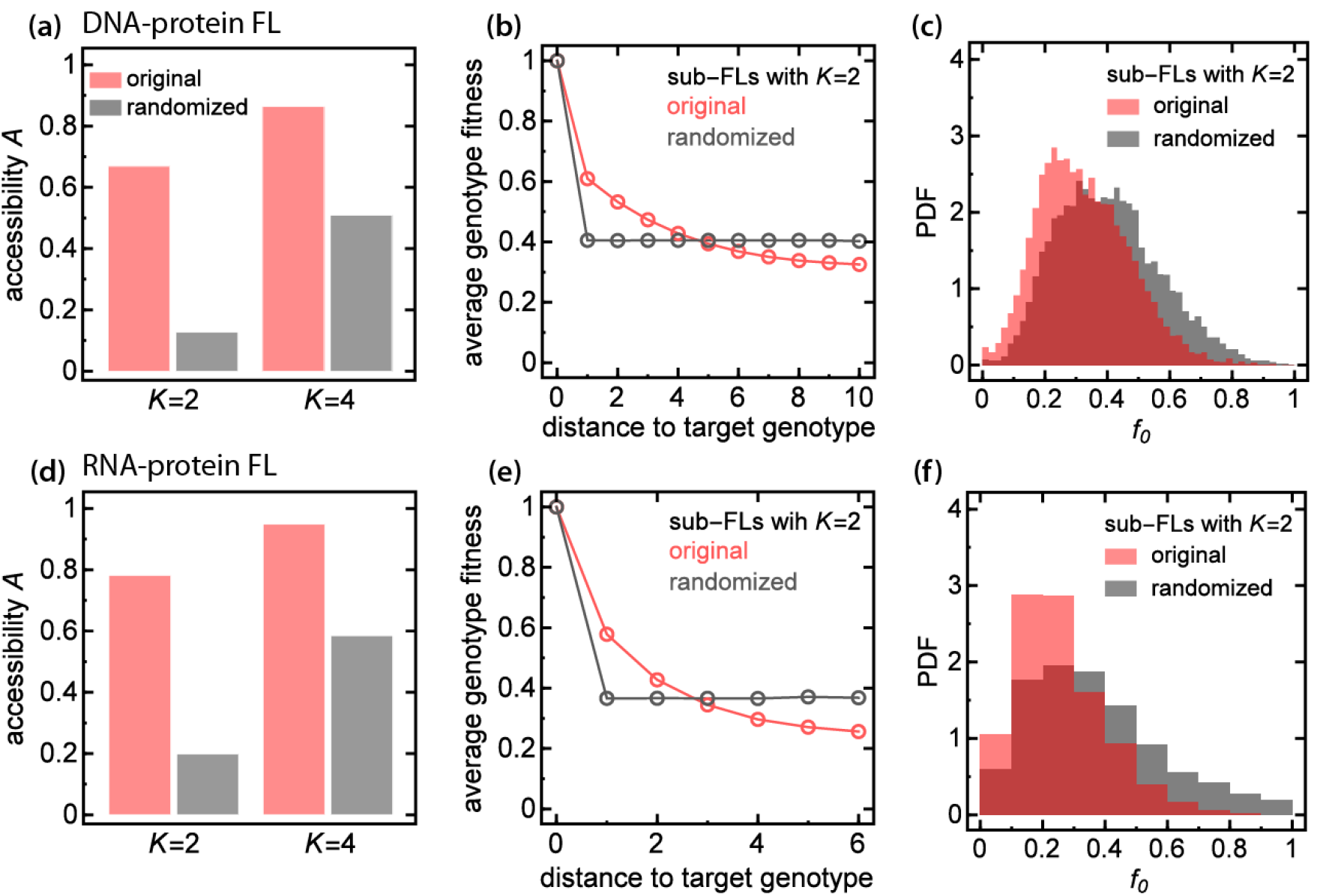
Accesibility of experimental FLs. Panels **(a-c)**: data from Rowe et al. [31], panels (d-f): data from Guenther et al. [33]. (a,d) Accessibility of the sub-FLs for *K* = 2,4 (red), and for their randomized counterparts with the same fitness distribution (grey). Randomization decreases accessibility to that of a maximally-random FL. **(b,e)** Average genotype fitness in sub-landscapes with *K* = 2 as a function of the distance from the target genotype. Randomization removes correlations present in the original FL. **(c,f)** Histogram of the fitness of the initial genotype; the average fitness for each histograms is the same as the average fitness value of the antipodal genotype (the right-most points in Panels b,e).

We further investigated how the fitness of the initial genotype affects accessibility, i.e., whether there is some critical value of initial fitness separating regions of FLs with high and low accessibility. To account for the non-uniform distribution of the experimental fitnesses we transformed the fitnesses such that the new fitness distribution was uniform, while preserving the order of the fitnesses and local correlations (epistasis), i.e., if *f*_0_ < *f*_1_ < *f*_2_ < … < *f_N–1_* then the transformed fitnesses 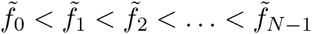. The estimated critical values of 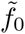 are 0.69(1) and 0.90(1) for *K* = 2 and *K* = 4 respectively (Fig 7a). These values are close to the estimated values of A from Fig. 6a; the critical *f_crit_* increases with *K* similarly as in the maximally random model.

**Fig 7.**
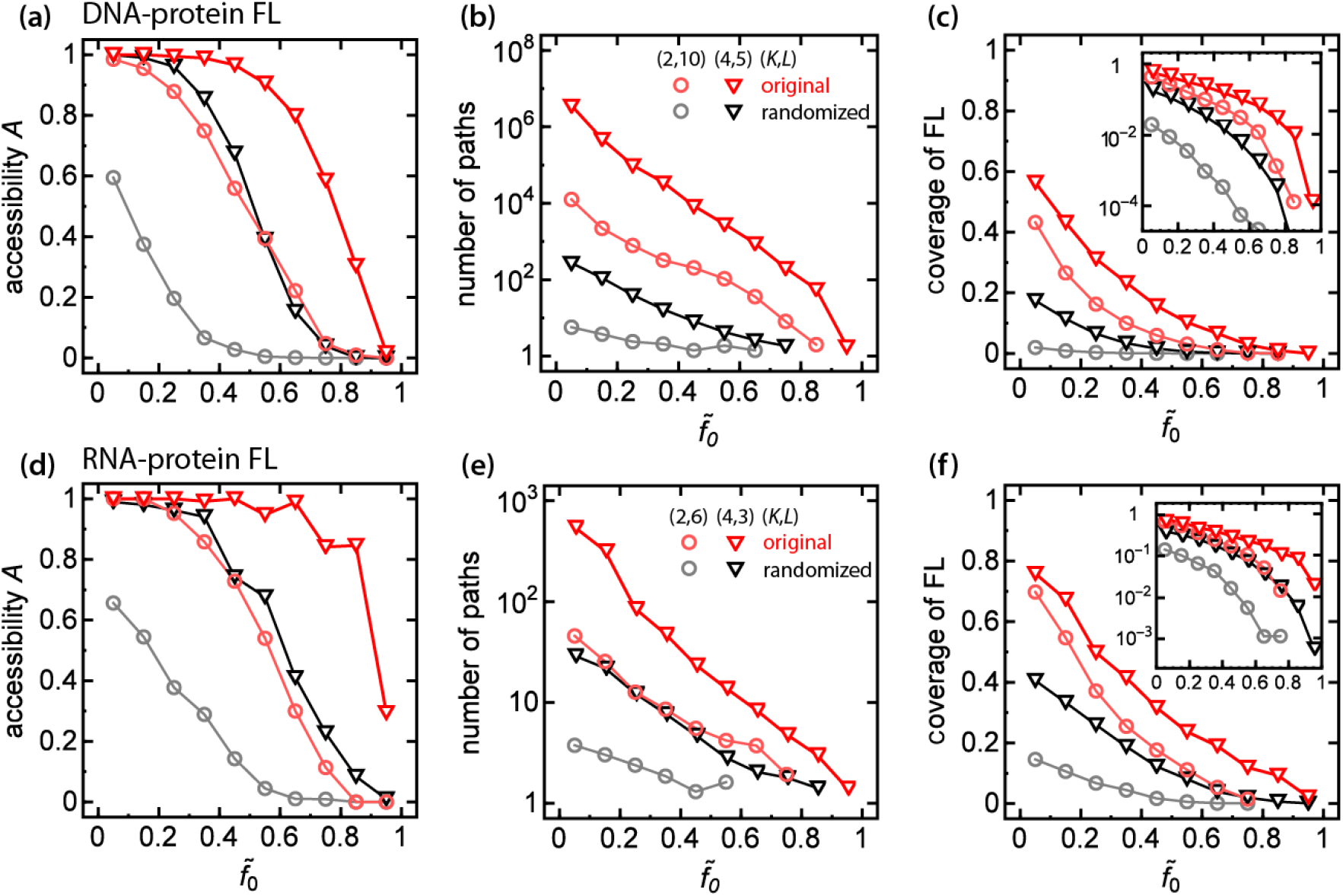
Fitness-to-distance correlation impacts accessibility and coverage of the experimental FLs. Panels **(a-c)**: data from Rowe et al. [31], panels (d-f): data from Guenther et al. [33]. **(a,d)** Accessibility of the sub-FLs for *K* = 2,4 (red circles, red triangles), and for their randomized counterparts (grey, black) as a function of rescaled initial fitness 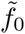. In the presence of fitness-to-distance correlations landscapes with high levels of 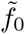 are mostly accessible compared to the case with no correlations. **(b,e)** Number of paths exhibits as a function of rescaled initial fitness 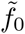. Presence of fitness-to-distance correlations increased number of pathways by a few orders of magnitude compared with the randomized ensemble. **(c,f)** Coverage of experimental sub-FLs and their randomized counterparts as a function of 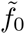. In the case of *K* = 2 correlations increase the fraction of accessible genotypes to levels observed for FLs with *K* = 4.

The presence of fitness-to-distance correlations affects also other properties of FLs: there are more accessible pathways leading to the best-fit genotype (Fig. 7b), the pathways are longer and the fraction of accessible genotypes is higher than in the maximally random model (Table S1). Moreover, the fractions of accessible genotypes for *K* = 2 and *K* = 4 are large and behave similarly when plotted against 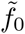 (Fig. 7c), but when fitness-to-distance correlations are removed the two curves separate. Correlations thus enhance accessibility for both *K* = 2 and *K* = 4, making *a posteriori* predictability of evolutionary trajectories even more difficult than in the case of the maximally-random FL.

In order to confirm that the observed behaviour is not unique to the specific experimental data set of Ref. [31], we analysed a second landscape of RNA-protein binding affinity [33]. We assumed fitness to be proportional to the rate constant for the reaction of the RNA-binding subunit C5 of RNase P with all variants of the 6-nucleotide recognition site (Methods). Proceeding similarly as before, we generated sub-FLs with *L* = 3, *K* = 4 and *L* = 6, *K* = 2 (*N* = 64 genotypes). The results for this second landscape (Fig. 6d-f, Fig. 7d-f, Table S1) turned out to be consistent with the results reported above.

## III. DISCUSSION

Although the concept of the fitness landscape was proposed almost a century ago, many open problems remain. Setting aside the appropriateness of the FL to model evolution in the presence of interactions between different organisms, one of the important questions is the accessibility of local fitness peaks: are such peaks sufficiently well connected, or are they separated by “fitness valleys” which are so deep that evolution cannot reach most of them? Another question concerns the predictability of evolution: how many evolutionary trajectories lead from one genotype to another, better adapted genotype?

Our results show that the answer to these questions depends on how genotypes are connected in the genotype space, i.e., which transitions between the genotypes are permitted and which are not. Since genetic information is stored in a linear molecule (DNA), genotypes are most naturally represented as linear sequences of some “units” of information, or building blocks, and this already imposes restrictions on the structure of the mutational graph. Moreover, the nature of the smallest unit of information and how mutations convert one unit into another further affects how genotypes are connected and how they are mapped to fitnesses. The choice of the appropriate “unit” depends on the subset of the fitness landscape considered in a particular research problem. While the presence or absence (K = 2) of a certain point mutation leads to a simple mathematical model, this approach is limited to small fragments of fitness landscapes and it requires the knowledge of which mutations are important and which can be ignored [10]. How many unit types are necessary to model the most general situation? Since the DNA code uses four nucleotides, *K* = 4 building blocks should in theory suffice to represent all possible genotypes. On the other hand, the redundancy of the genetic code might suggest that the most appropriate unit should rather be an amino acid, in which case *K* ≈ 20. However, this could be disputed because there is evidence that synonymous codons can be translated at different rates [34–36] which may affect fitness.

We have seen that increasing the number of basic units or “alleles” *K* generally increases accessibility. This is not just a trivial consequence of increased connectivity of the mutational graph (number of permitted mutations for each genotype). Connectivity also increases with increasing sequence length *L*, but this actually slightly decreases accessibility (cf. Fig. 2). Increasing *K* changes not only how many neighbours a genotype has in the genotype space, but it affects the global topology of the network of genotypes. Since accessibility of the global fitness maximum is a global property of the fitness landscape, it depends on *K*. This is in contrast to local properties such as the number of local fitness maxima, which depend only on the number of nearest neighbours and is thus the same for FLs with the same value of the product (*K* – 1)*L* (Fig. S5).

The presence of multiple alleles has also one more important effect - it enables pathways to explore a much larger part of the fitness landscape through mutations that either do not decrease the Hamming distance to the target genotype or even increase it (backward mutations). This greatly enhances accessibility by exploring genotypes through indirect mutations, effectively circumventing regions of sign epistasis [37–39]. This observation has been recently supported by experimental study of the four-site IgG-binding domain of protein *G* (*L* = 4, *K* = 20) [40] which shows that direct paths are often blocked by reciprocal sign epistasis, and only indirect pathways can access high-fitness regions of the landscape.

By re-sampling two experimental fitness landscapes [31, 33] we have found that the presence of correlations between the fitnesses facilitates evolution and, as a result, accessibility and the fraction of accessible genotypes for *K* = 2 increases to a level comparable with that of the maximally-random FL with *K* > 2. For larger *K* the increase of accessibility is not so pronounced (accessibility is already high in the maximally random FL), but the number of accessible pathways increases by many orders of magnitudes: there are ~ 10^10^ pathways in the presence of correlations as compared to ~ 10^3^ pathways in the randomized FL with 1024 genotypes (Table S1). It would be interesting to see if similar effects could be observed in other large FLs [41, 42]. However, these data sets, despite a large number of genotypes included, cover only a small fraction of the genotype space due to random mutagenesis employed to generate mutant genotypes. The problem of missing genotypes with undefined fitness is why in this work we decided to use only the (almost complete) FLs of Refs. [31, 33].

Our analysis, carried under the assumption of low mutation probability and strong selection, assumes that all evolutionary pathways are equally likely. This does not have to be true in many biologically relevant scenarios [2, 43, 44]. It has also been shown that pathways with backward mutations have little statistical weight when *K* = 2 [45]. Our work shows that such pathways are anyway rare for *K* = 2, but they dominate the set of accessible pathways when *K* > 2. It may therefore happen that once the low mutation/strong selection assumption is relaxed, indirect pathways become less common.

Another factor that we neglected was that genotype-to-phenotype mapping is not unique due to phenotypic plasticity [46–48] in which many phenotypes may correspond to the same genotype. Moreover, fitness is not an absolute characteristic of an organism in the same way as the genotype is because it depends on the organism’s environment: physical conditions such as temperature, available nutrients, and the presence/absence of other organisms. This environmental plasticity has been shown to speed up evolution in experiments [49–51] and *in silico* [52–54]. Fitness can also be different at different locations even in a relatively small habitat, and chemical gradients can significantly affect the rate of biological evolution [55, 56]. The reality is thus much more complicated than our idealised model, but the picture that emerges from our work is that the mutational structure of the genotype space is an important factor affecting biological evolution, and that the use of binary sequences to represent real genotypes has limited applicability.

## IV. METHODS

### Computer model

For each generated fitness landscape (FL) we find the number of accessible pathways that start from the antipodal genotype {0,0, …0} and end at the fittest (target) genotype {*K* – 1, *K* – 1, …, *K* – 1} using a depth-first search (DFS) algorithm with backtracking. The fitness landscape is represented as a graph with nodes corresponding to genotypes and edges to mutations between the pairs of genotypes. Each node is assigned a counter variable which specifies how many accessible pathways connect this node to the fittest genotype, and an auxiliary variable *υ* which assumes one of the three possible node states: *υ* = 0, *υ* = 1 or *υ* = 2, representing nodes that have not been visited by DFS yet, the visited ones, and the ones with their whole mutational neighbourhood explored. All counters and variables v are initially set to zero. The DFS algorithm starts from the antipodal genotype and follows only those edges along which fitness increases. When the final node (genotype with maximal fitness) has been reached the algorithm traces its steps back to the initial node while increasing the counters of all nodes it visits on its way. Particularly, if all edges from a given node have been explored, the node is labelled as explored (*υ* = 2) and its counter variable is no longer updated. Next, the counter of this node is added to the counter of the previous node in the DFS and the search continues from the later node. If at some point the DFS algorithm attempts to visit an already explored node, only the explored node counter is added to the counter of the current node. The backtracking procedure thus exhaustively accumulates the number of all paths going from a given node to the target genotype. When the search is completed, the value of the counter at the initial node gives the total number of accessible paths starting at this node.

We tested this algorithm by comparing the number of pathways found for small (*N* < 2^10^) FLs with the number obtained by a naive, recursive breath-first search algorithm. Both algorithms produced identical results but the first algorithm was substantially faster for larger graphs than the naive algorithm. For example, the described algorithm enabled us to enumerate all pathways (~ 10^34^) on a computer-generated FL with *K* = 16, *L* = 6 which would clearly be impossible for the naive algorithm.

To obtain the accessibility of a given FL we used a simple DFS algorithm without backtracking. This reduced memory requirements as compared to the full algorithm described above, enabling FLs with higher *K* and/or *L* to be analysed. To obtain the number of forward, backward and distance-neutral mutations in pathways of a given length, we modified the algorithm so that it stored a two-dimensional histogram – a generalized counter variable – for each node. During backtracking the histograms were accumulated and stored as the final histogram in the antipodal genotype (in Fig. S6 example of such histogram is plotted for the experimental FL with *K* = 4 and *L* = 10).

### Experimental fitness landscape

The first experimental FL we used was the protein-binding landscape of DNA oligomers of length *L* = 10 [31]. This landscape has been previously shown to be rugged, with many minima and maxima, and is thus not too dissimilar from our computer-modelled landscapes. However, the landscape shows strong correlations (Fig. 2 in Ref. [31]) and thus it is not evident how our results from the computer model of maximally-random, uncorrelated FLs will apply to this landscape.

The natural number of coding units, or “alleles”, for this landscape is *K* = 4 since each site of the sequence can be in one of four possible states (4 nucleotides A,T,G,C). The total number of genotypes is 4^10^ ≈ 1M. In order to obtain quasi-independent fitness landscapes necessary for statistical analysis from just one, large data set and FLs for other values of *K* (in particular for *K* = 2), we followed the procedure described below.

To obtain a sample landscape with *K* = 4, *L* = 5, we randomly selected 5 different positions 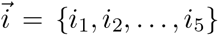 from the 10-nucleotide sequence, and a random sequence 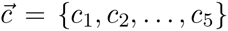 where *c_i_* was one of A,T,G, or C. We then selected all 10-nucleotide sequences with nucleotides 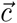 at positions 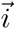, and any nucleotides at the remaining *L* = 5 positions. This generated a sub-landscape of the full fitness landscape with *N* = 4^5^ = 1024 genotypes with the same “genetic background” at positions 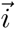. By repeating the above procedure we created 10k (of the total ≈ 258k different) FLs with different genetic backgrounds (different 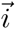).

To generate a sample landscape with *K* = 2, *L* = 10, we selected a random sequence 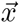 from all 10-nucleotide sequences, and another sequence 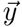 such that it differed from 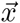 at all positions. We associated sequences 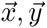 with two antipodal binary sequences 00… 0 and 11… 1 of length *L* = 10, and constructed the fitness landscape from all *N* = 2^*L*^ = 1024 sequences such that the *i*-th nucleotide was either *x_i_* or *y_i_*. We repeated this procedure to sample 10k landscapes of the total ≈ 60M different possible FLs.

After generating the sets of fitness landscapes with *K* = 4 and *K* = 2 with the same number of genotypes *N* = 1024, the corresponding fitness values were normalized to be between 0 and 1. Accessibility was determined for mutational pathways starting at the antipodal genotype and ending at the genotype with maximum fitness. For *K* = 4 we used the same convention of selecting the antipodal genotype as in the computer-generated FLs (see below).

For our second FL we used the rate constants for the reaction between the RNA-binding subunit C5 of RNase P recognition and its RNA substrate, with all variants of its recognition site of length *L* = 6 [33]. Using the same approach as described in the Methods of Ref. [33] we calculated the rates 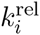 relative to the rate constant of the wild-type RNA sequence. In our analysis we were less stringent than Ref. [33] on removing sequence variants that exhibited biased amplification, i.e., we kept sequences that had reads above quality threshold for at least three time points. We took as fitness the relative rate constant averaged over different time points; we also checked that other methods, e.g. selecting 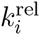 with the largest difference in the corresponding raw read count, did not significantly affect our results. The analysis yielded fitness values for 3940 out of the total 4^6^ = 4096 possible sequences. The remaining 4% sequences for which fitness could not be determined were assigned the fitness of the least fit genotype.

Adapting the procedure described for the first experimental landscape we obtained sublandscapes of the full experimental landscape with *K* = 4, *L* = 3 and *K* = 2, *L* = 6, with *N* = 64 genotypes each. Since these sub-FLs were sampled from the landscape smaller than the first one and the total number of possible sub-FLs was much smaller than before (e.g., 1280 for *K* = 4), we generated only 1k sub-FLs for each *K*. We also normalized the fitness to be between 0 and 1, and performed the same analysis as for the first experimental data set.

### Re-indexing fitness landscapes

In both the computer model and the analysis of the real FLs we re-index all genotypes such that the best-fit (target) genotype is assigned the sequence {*K* – 1, *K* – 1, …, *K* – 1}. The antipodal genotype is then {0,0, …0}. When fitness values are drawn at random no actual re-indexing is necessary but the series of *N* = *K^L^* fitnesses is generated and the highest value is assigned to the target genotype while the remaining fitness values are assigned to the remaining genotypes. For sub-landscapes of the real fitness landscapes we define a map that assigns the target genotype sequence to the fittest genotype while preserving all correlations and the structure of the FL; in particular, each genotype has the same nearest neighbours in the re-indexed FL. For *K* = 2 we proceed as follows: if the fittest genotype sequence has 0s at *k* positions 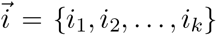 and 1s at the remaining positions, the map changes 0 → 1 and 1 → 0 at the 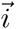 positions and for the remaining positions there are no changes. Applying this transformation to all genotypes results in a FL with the desired indexing (the fittest genotype sequence contains only 1s and the antipodal genotype sequence only 0s). This mapping is unique for *K* = 2, but in the case of *K* > 2 the antipodal genotype can be selected in many ways. We thus modify the algorithm as follows for *K* > 2. Assume that the fittest genotype sequence has characters 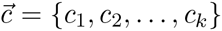 at positions 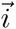 such that 0 ≤ *c*_1_, *c*_2_, …, *c_k_* < *K* – 1. We then define the following reassignment: *c*_1_ → *K* – 1, …, *c_k_* → *K* – 1, which maps the fittest genotype to the sequence {*K* – 1, *K* – 1, …, *K* – 1}. There are now (*K* – 1)^*L*^ possible choices for the antipodal genotype and we remove this ambiguity by randomly selecting one of them and extending the mapping (character reassignment) in such a way that the selected genotype is mapped onto the sequence {0,0, …0}. The character reassignment rules described above are then applied to all remaining genotypes. We note that these rules apply only to specific positions in the sequence; for example the mapping 1 → *K* – 1 at position 3 will be applied only if a sequence has character 1 at position 3, but not if there is a 1 at another position, unless there is a rule 1 → *K* – 1 specifically for that position. We also note that the algorithm does not change characters which are neither assigned to (*K* – 1) (target) nor to 0 (antipodal).

## ACKNOWLEDGMENTS

We thank Marjon de Vos and Oliver Martin for critically reading the manuscript.

## AUTHOR CONTRIBUTIONS

MZ, ZB, BW conceived and designed the research. MZ, BW wrote the computer programs, performed the simulations and analysed the data. MZ, ZB, BW discussed the results and wrote the manuscript.

## Supporting Information

**Fig S1.**
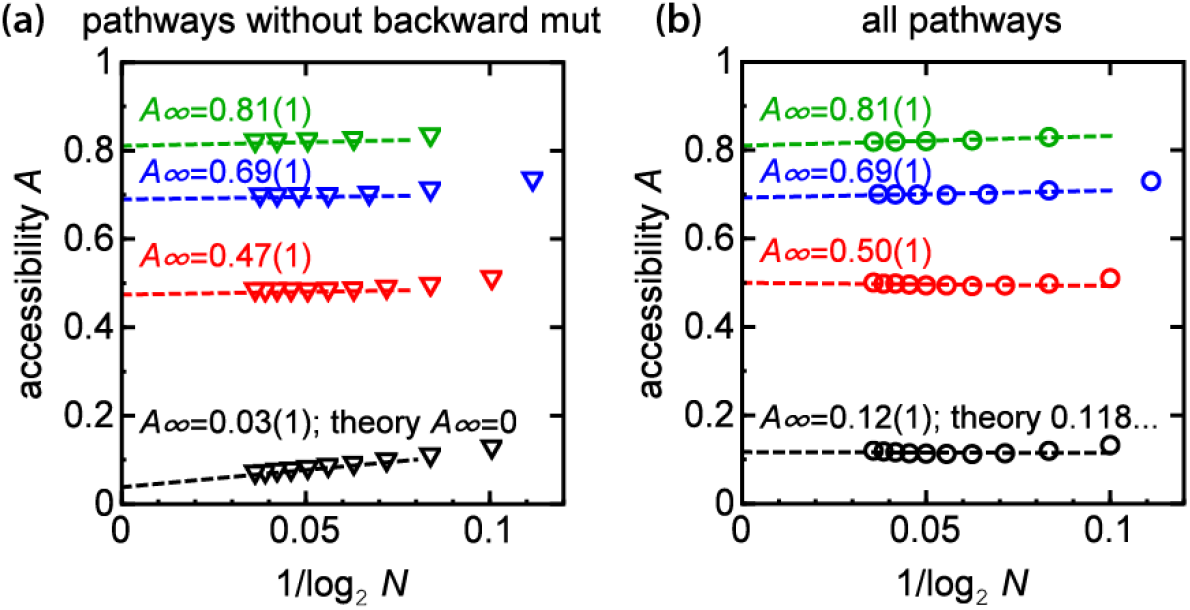
Asymptotic accessibility estimates for different *K*. Plots of accessibility *A* versus the inverse of log_2_ *N* for different *K* (2=black, 4=red, 8=blue, and 16=green) and for pathways without backward mutations (panel a) and for all pathways (panel b). The dashed lines are linear fits to the data.

**Fig S2.**
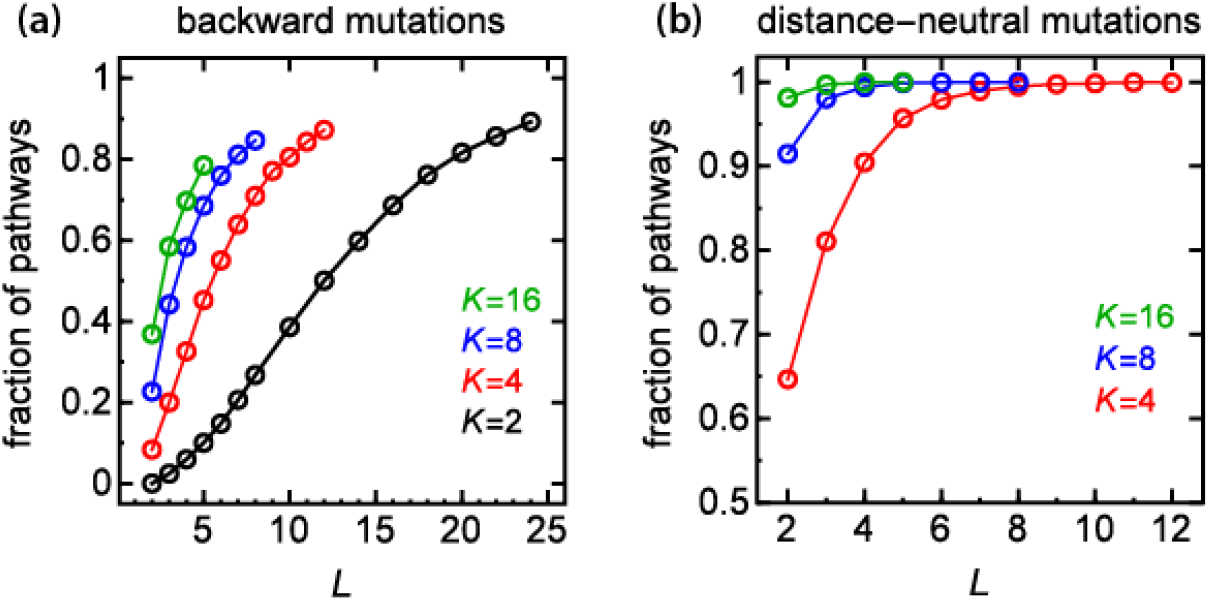
Fraction of pathways with backward and distance-neutral mutations increases with *K*. **(a)** Average fraction of pathways that contain at least one backward mutation as a function of genotype length *L*, and for different *K*. **(b)** Analogous to panel a, but for distance-neutral mutations. By definition, distance-neutral mutations do not exist for *K* = 2 and hence there is no corresponding line in the plot.

**Fig S3.**
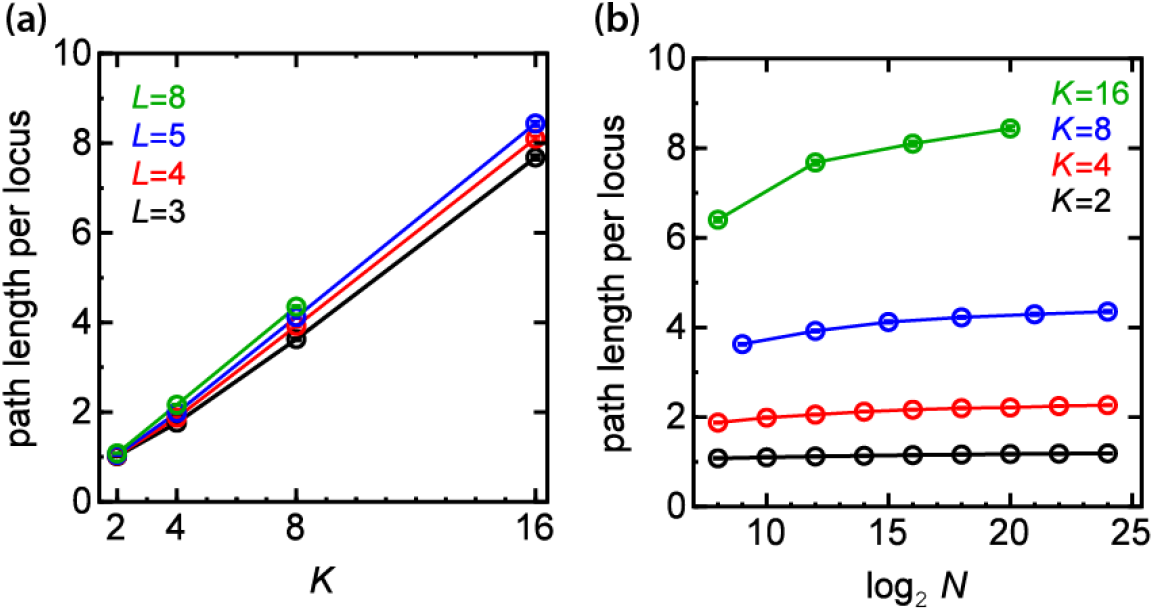
Increasing the number of coding units lengthens accessible pathways. **(a)** The normalized pathway length (number of mutational steps divided by *L*) as a function of *K*, for different values of *L*. **(b)** The normalized length as a function of log_2_ *N* for different values of *K*. Except from *K* =16 the normalized length reaches a plateau within the investigated range of *N*.

**Fig S4.**
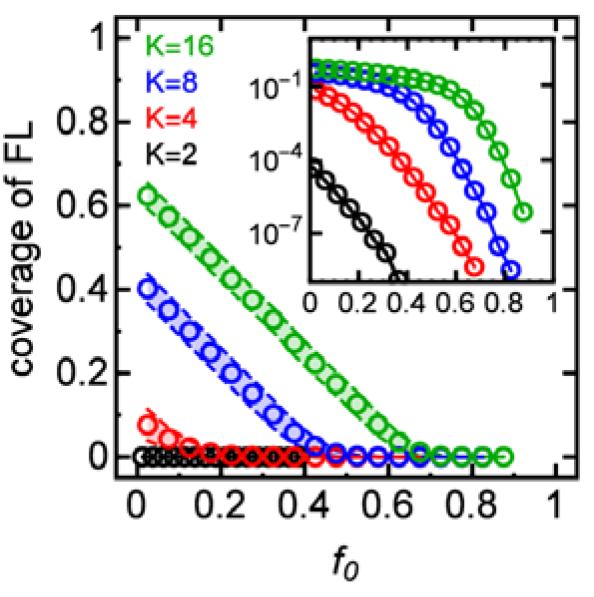
Coverage of FL with genotypes belonging to accessible pathways depends on the fitness of the initial genotype. Average coverage of fitness landscapes with different number of coding units *K* (black, red, blue, green) and a fixed number of genotypes (*N* = 2^24^ for *K* = 2,4,8 and *N* = 2^20^ for *K* = 16) as a function of initial fitness *f*_0_. Inset: the same data in the log-linear scale. Shaded area corresponds to one standard deviation (accessible FLs only).

**Fig S5.**
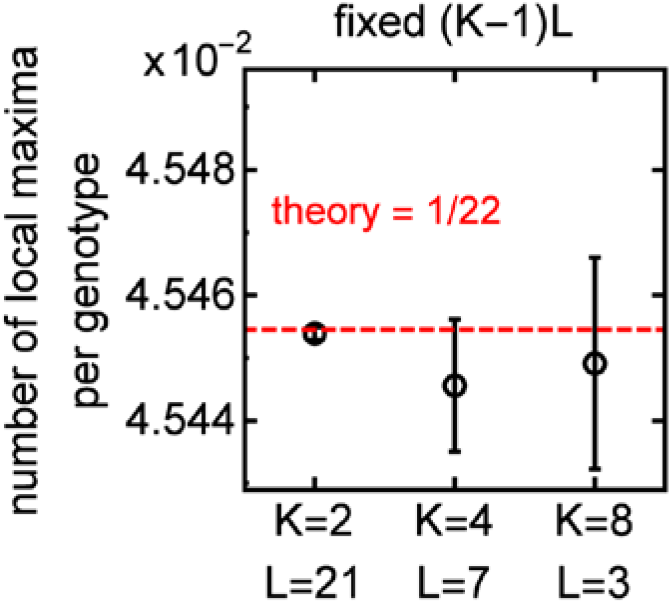
Number of local fitness maxima depends only on the local connectivity of FL. Number of local fitness maxima in random fitness landscapes with different *K* is determined by local properties of the FL (the number of neighbours (*K* – 1)*L* in the genotype space). This is not the case for accessibility which depends on the global structure of FL.

**Fig S6.**
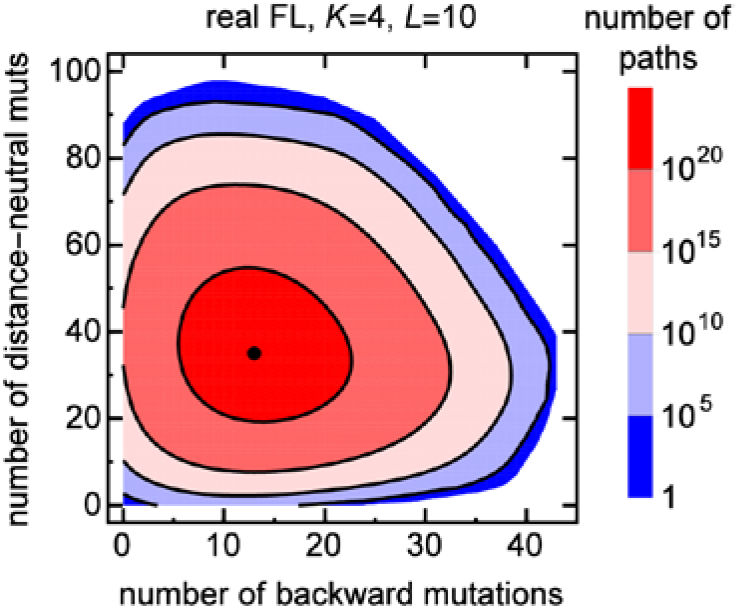
Number of accessible pathways in the experimental FL has a broad distribution in the space of indirect mutations. Number of accessible pathways from the antipodal to the best-fit genotype for the full experimental FL as a function of the number of backward and distance-neutral mutations. The maximal number of pathways (black dot) is approximately 6.45 × 10^21^. The histogram was obtained by exhaustive enumeration of all pathways using the algorithm described in Methods.

**Tab S1.** Fitness-to-distance correlations facilitate accessibility of the best-fit genotype. Properties of the two fitness landscapes generated from the experimental data sets discussed in the main text. Randomized ensembles of FLs preserve fitness values, but not fitness-to-distance correlations.

**10k sub-FLs generated from DNA-protein binding affinities (Rowe et al.)**
| | $K=4, L=5$ | | $K=2, L=10$ | |
| --- | --- | --- | --- | --- |
|  | Experimental | Randomized | Experimental | Randomized |
| accessibility $A$ | 0.86 | 0.51 | 0.67 | 0.13 |
| $\langle \text{coverage of FL} \rangle^*$ | 28% | 8% | 26% | 3% |
| $\langle \text{pathway length} \rangle$ | 16.8 | 10.0 | 15.4 | 11.1 |
| $\langle \text{forward mutations} \rangle$ | 43% | 57% | 82% | 95% |
| $\langle \text{distance – neutral mutations} \rangle$ | 44% | 37% | 0% | 0% |
| $\langle \text{backward mutations} \rangle$ | 13% | 7% | 18% | 5% |
| $\langle \text{pathway number} \rangle$ | $1.2 \times 10^{10}$ | $1.9 \times 10^3$ | $2.3 \times 10^6$ | 19 |
| $\langle \log_{10}(\text{pathway number}) \rangle$ | 5.4 | 2.1 | 3.9 | 0.8 |

**1k sub-FLs generated from RNA-protein binding affinities (Guenther et al.)**
| | $K=4, L=3$ | | $K=2, L=6$ | |
| --- | --- | --- | --- | --- |
|  | Experimental | Randomized | Experimental | Randomized |
| accessibility $A$ | 0.95 | 0.59 | 0.78 | 0.20 |
| $\langle \text{coverage of FL} \rangle$ | 42% | 24% | 44% | 19% |
| $\langle \text{pathway length} \rangle$ | 6.7 | 5.3 | 7.1 | 6.4 |
| $\langle \text{forward mutations} \rangle$ | 52% | 61% | 92% | 97% |
| $\langle \text{distance – neutral mutations} \rangle$ | 41% | 35% | 0% | 0% |
| $\langle \text{backward mutations} \rangle$ | 7% | 4% | 8% | 3% |
| $\langle \text{pathway number} \rangle$ | $1.8 \times 10^3$ | 58 | 131 | 6.1 |
| $\langle \log_{10}(\text{pathway number}) \rangle$ | 2.2 | 1.2 | 1.6 | 0.6 |

<sup>\*</sup>) The average $\langle \dots \rangle$ was restricted to accessible FLs: for each such FL we first calculated the corresponding quantity, and then averaged it over all accessible FLs. Statistical errors (s.e.m.) are at most +/-1 of the last digit.

